# 12-Lipoxygenase Governs the Innate Immune Pathogenesis of Islet Inflammation and Autoimmune Diabetes

**DOI:** 10.1101/2021.01.02.424855

**Authors:** Abhishek Kulkarni, Annie R. Pineros, Sara Ibrahim, Marimar Hernandez-Perez, Kara S. Orr, Lindsey Glenn, Melissa Walsh, Jerry L. Nadler, Margaret A. Morris, Sarah A. Tersey, Raghavendra G. Mirmira, Ryan M. Anderson

## Abstract

Macrophages and related myeloid cells are innate immune cells that participate in the early islet inflammation of type 1 diabetes (T1D). The inflammatory signals and antigen presentation by these cells may be inducers of the adaptive immune response that is the hallmark of T1D. The enzyme 12-lipoxygenase (12-LOX) catalyzes the formation of pro-inflammatory eicosanoids from membrane-derived phospholipids, but its role and mechanisms in the pathogenesis of islet inflammation have not been elucidated. Leveraging a model of T1D-like islet inflammation in zebrafish, we show here that macrophages contribute significantly to the loss of β-cells and the subsequent development of hyperglycemia. Depletion or inhibition of 12-LOX in this model resulted in significantly reduced macrophage infiltration into islets with preservation of β-cell mass. In mice, we deleted the gene encoding 12-LOX (*Alox15*) in the myeloid lineage in the non-obese diabetic (NOD) model of T1D. Myeloid cell-specific *Alox15* knockout NOD mice demonstrated reduced insulitis and T-cell responses, preserved β cell mass, and almost complete protection from the development of T1D. A critical effect of 12-LOX depletion appeared secondary to a defect in myeloid cell migration, a function required for immune surveillance and tissue injury responses. This effect on migration appeared to be secondary to the loss of the chemokine receptor CXCR3. Transgenic expression of the gene encoding CXCR3 rescued the migrator defect in zebrafish 12-LOX morphants. Taken together, our results reveal a formative role for innate immune myeloid cells in the early pathogenesis of T1D and identify 12-LOX as a necessary enzyme to promote their pro-diabetogenic phenotype in the context of autoimmunity.

## INTRODUCTION

The two major forms of diabetes, type 1 (T1D) and type 2 (T2D), represent disorders of glucose homeostasis whose common feature is the failure of the islet β cell to secrete adequate insulin [1]. The underlying etiologies of β-cell failure in each form of diabetes differs. In the case of T1D, an active dialog between β cells and cells of the immune system results in the cytokine-induced dysfunction of β cells in the early phase of disease, and in the destruction of β-cells by autoreactive T-cells in the later phase [2]. A pathology that characterizes this dialog is “insulitis,” a feature in which islets are invaded by cells of the myeloid and lymphoid lineage [3]. Insulitis and the nature and timing of cells invading the islet have been characterized best in the non-obese diabetic (NOD) mouse model of T1D [3,4]. Insulitis has also been demonstrated in human T1D, albeit at a much lower frequency than that in NOD mice [5]. Drug targeting of the immune system has been the foundational approach in clinical studies attempting to prevent or reverse T1D, but these studies have targeted the activation or function of lymphoid cell responses with variable and transient effects [6].

An emerging perspective in T1D posits that islet inflammation arising from innate immune responses may initiate the formation of neoepitopes in β cells, thereby signaling a cascade of immune signaling events leading to the development of T cell-mediated autoimmunity [7]. Central to this perspective are cells of the myeloid lineage, which include macrophages and dendritic cells. These cells mediate inflammation and present antigen to the adaptive immune system [8], and have been observed in the early stages of insulitis in mouse models of T1D [9–11]. The presence of “resident” macrophages and their closely-related antigen-presenting dendritic cells have been demonstrated in early insulitic lesions in NOD mice [9,12], and their functional inhibition via antibody- or clodronate-mediated global sequestration slows or prevents the occurrence of T1D [11,13–16]. However, as a strategy for T1D, the therapeutic depletion of macrophages would be undesirable given the consequences on overall immunity. Nevertheless, the identification of amenable targets that function primarily in myeloid cells during the pathogenesis of T1D would complement existing lymphoid cell-targeting strategies.

Lipoxygenases (LOX) are enzymes that catalyze the di-oxygenation of polyunsaturated fatty acids. Specifically, 12/15-lipoxygenase (referred to henceforth as simply 12-LOX), which is predominantly expressed in macrophages and pancreatic islets in mice [17], catalyzes the conversion of arachidonic acid to the eicosanoids 12- and 15-hydroxyeicosatetraenoic acid (12- and 15-HETE). Global deletion of the gene encoding 12-LOX in mice (*Alox15*) on the NOD background results in near-total protection of both sexes from T1D development, with a striking reduction in insulitis and the early accumulation of macrophages [18]; similarly, delivery of a small molecule inhibitor of 12-LOX (ML351) in NOD mice shortly after the development of insulitis protects against progression of insulitis and glycemic deterioration [19]. Whereas these findings provide evidence for the safety and efficacy of targeting 12-LOX in the context of T1D, they leave unclarified the specific cell of action and molecular mechanisms of 12-LOX. In this study, we leveraged genetic models in zebrafish and mice to investigate the role of 12-LOX in the myeloid cell pathogenesis of T1D. Our findings provide evidence for a determinative role for myeloid 12-LOX in the initiation of T1D and highlight the seminal role of innate immunity in the propagation of T1D autoimmunity.

## RESULTS

### Development of a zebrafish platform to interrogate the role of macrophages in the early pathogenesis of T1D

The zebrafish is a powerful model organism to test hypotheses and mechanisms that can then be interrogated further in mammalian systems. Developing zebrafish (3-4 days postfertilization, dpf) exhibit a discrete pancreatic islet with functional β-cells [20], and intact innate immunity with *mpeg*-expressing macrophages [21]. We utilized a model of chemically-induced β-cell oxidative injury that we described previously [22], in which zebrafish harboring β-cell-specific expression of the gene encoding bacterial nitroreductase (NTR) are treated with the prodrug metronidazole (MTZ). We crossed transgenic β-cell NTR-expressing zebrafish (*Tg(ins:NTR)*) with transgenic fish containing enhanced green fluorescent protein (eGFP)-labeled macrophages (*Tg(mpeg:eGFP)*) to generate a double-transgenic fish line that allows for chemical β-cell injury and simultaneous visualization of macrophages in real-time. Upon treatment of double-transgenic 3 dpf zebrafish with 7.5 mM MTZ (Fig. 1A), we observed rapid loss (within 12 hours) of β cells with influx of macrophages and evidence of β-cell phagocytosis (Fig. 1B). Glucose levels increased within 24 hours following MTZ treatment (Fig. 1C, blue line) compared to untreated control fish, whose levels remained unchanged (Fig. 1C, black line). To demonstrate that β-cell damage and loss were at least partially attributable to macrophage activity, we repeated the MTZ ablation experiment following trans-pericardial injection with 5 mg/mL clodronate liposomes at 3 days post-fertilization. Clodronate sequesters and depletes macrophages [23]. In clodronate-injected fish we observed significant reduction in hyperglycemia (Fig. 1C, red line) and preservation of β-cell number compared to control-injected fish (Fig. 1D-E). These data suggest that our MTZ model in zebrafish mimics the early phases of T1D where inflammation driven by macrophages contributes to the development of hyperglycemia [13,14].

**Figure 1:**
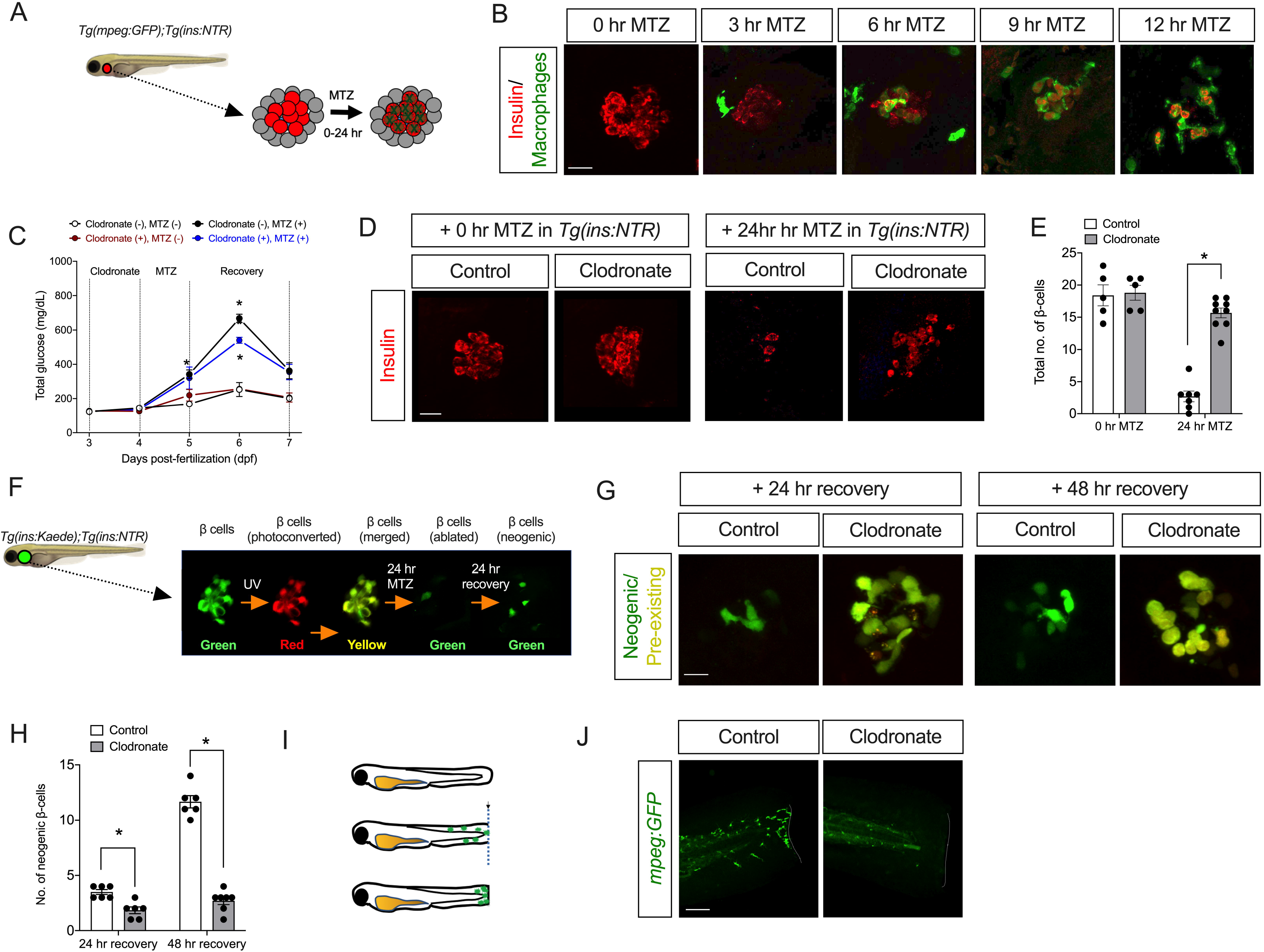
Macrophages promote β-cell loss and hyperglycemia following injury in zebrafish. (***A***) Schematic representation of the β-cell injury assay in transgenic *Tg(mpeg:GFP);Tg(ins:NTR)* zebrafish at 3 dpf, wherein islet β cells (*red*) are selectively destroyed upon incubation with metronidazole (*MTZ*), with the concomitant/subsequent entry of macrophages (*green*); (***B***) Representative images of islets from zebrafish treated for the times indicated with MTZ, then stained for insulin (*β cells, red*) and GFP (*macrophages, green*). Scale bar indicates 10□μm; (***C***) Free glucose measurements of whole zebrafish lysates, treated as indicated in the panel. N=3-5 lysates per condition (20 fish per lysate), and \**P*<0.05 for the corresponding values compared to untreated controls (no clodronate, no MTZ); (***D***) Representative images of islets from zebrafish stained for insulin (*β cells, red*) under the conditions indicated. Scale bar indicates 10□μm: (***E***) Quantitation of β-cell number from the experiment represented in *panel **D***; (***F***) Schematic representation of the β-cell regeneration assay, where photoconversion of *Kaede* protein results in *red+green (=yellow*) pre-existing β cells and newly formed (neogenic) β cells enter as *green* cells. (***G***) Representative images of islets from zebrafish exhibiting pre-existing (*yellow*) and neogenic (*green*) β cells at 24 and 48 hr of recovery following MTZ treatment under the conditions (control or clodronate) indicated. Scale bar indicates 10□μm: (***H***) Quantitation of neogenic β cell number from the experiment represented in *panel **G***; (***I***) Schematic representation of the zebrafish tailfin injury assay, where tailfins of *Tg(mpeg:GFP)* fish at 3 dpf are mechanically cut with a blade, and the migration of macrophages (*green*) are observed at the site of injury; (***J***) Representative tailfin images of injured zebrafish tails stained with GFP (*macrophages, green*) under the conditions indicated. *Dotted line* shows the tailfin injury site. Scale bar indicates 100 μm. In all panels, data are presented as mean□±□SEM. \**P* <0.05 for the comparisons shown.

Because zebrafish β cells have a capacity for rapid regeneration, we asked if the apparent preservation of β-cell number upon depletion of macrophages was a reflection of the survival of β cells following MTZ treatment or their rapid replacement by neogenesis. To distinguish between these possibilities, we utilized the *Tg(ins:Kaede^s949^)* transgenic zebrafish line, which drives expression of a green photoconvertible protein and thus permits the pulse labeling of β cells with green and/or UV-converted red fluorescence (Fig. 1F). In this transgenic line, pre-existing β cells will be labeled yellow following photoconversion (combined red and green), whereas neogenic β-cells formed after the UV exposure will be labeled green (Fig. 1F). We generated double-transgenic (*Tg(ins:NTR);(ins:Kaede)*) zebrafish then treated them with MTZ to induce β-cell stress. When β cells were injured, the control-injected fish showed β-cell neogenesis at 24 hours of recovery that increased 3-fold by 48 hours of recovery (Fig. 1G-H). By contrast, at both 24 and 48 hours after β-cell ablation, clodronate-injected fish showed significantly decreased β cell neogenesis relative to controls, with no increase at 48 hours (Fig. 1G-H). These data indicate that β-cell number is greater in the fish lacking macrophages because of persistence of pre-existing β cells rather than β-cell neogenesis.

To ensure that our macrophage tissue injury model was not peculiar to MTZ treatment, we also performed mechanical tailfin injury assays in 3 dpf *Tg(mpeg:eGFP)* zebrafish. In the tailfin injury assay (shown schematically in Fig. 1I), macrophages rapidly migrate to the injury site as part of an inflammatory response [24]. Following the injury, macrophages were observed to migrate to the site of tissue injury, as expected, whereas in clodronate-injected fish macrophages were not observed at the site of injury (Fig. 1J). Taken together, the data in Fig. 1 support an inflammatory model in zebrafish, where β-cell injury is effected, in part, by macrophages and where macrophage depletion reduces β-cell loss and mitigates hyperglycemia.

### 12-LOX is required for macrophage-directed β-cell injury

Previous studies in mouse models of T1D demonstrated that global deletion or chemical inhibition of the enzyme 12-LOX reduces the early infiltration of macrophages into islets, preserves β-cell mass, and prevents hyperglycemia [18,19]. We recently demonstrated the presence of a zebrafish 12-LOX ortholog encoded by *alox12*, which exhibits similar catalytic activity and product profiles (including the production of 12-HETE) to the mouse and human enzymes [25]. To test a role for 12-LOX in β-cell injury mediated by macrophages, we utilized the zebrafish β-cell injury model in the transgenic line *Tg(ins:NTR)* following depletion of 12-LOX by injection of a translation-blocking antisense morpholino (*alox12* MO) [25]. With MTZ treatment there was a time-dependent reduction in β-cell number, with only 6% of β cells remaining at 24 h (Fig. 2A-B). By contrast, there was significant preservation of β cells when the fish were treated with *alox12* MO, with about 25% of β cells remaining at 24 h following MTZ (Fig. 2A-B). Next, we utilized the double-transgenic line *Tg(ins:NTR);(mpeg:eGFP)* to track the influx of macrophages following β cell injury in the presence or absence of *alox12* MO. As expected, MTZ treatment resulted in the migration of macrophages into the islet in control-injected fish (Fig. 2C-D), whereas in *alox12* MO fish there was a significant 2.7-fold reduction in macrophage numbers (Fig. 2C-D). The reduction of macrophages at the site of β-cell injuring following the *alox12* MO was likely due to a reduction of macrophage migration rather than reduction in total macrophage numbers, since macrophages persist elsewhere in the embryo (unlike with clodronate injection) (Supplementary Fig. S1A), and similar macrophage numbers were seen in the immediate vicinity of the islet in injured *alox12* MO fish compared to control-injected fish (Supplementary Fig. S1B).

**Figure 2:**
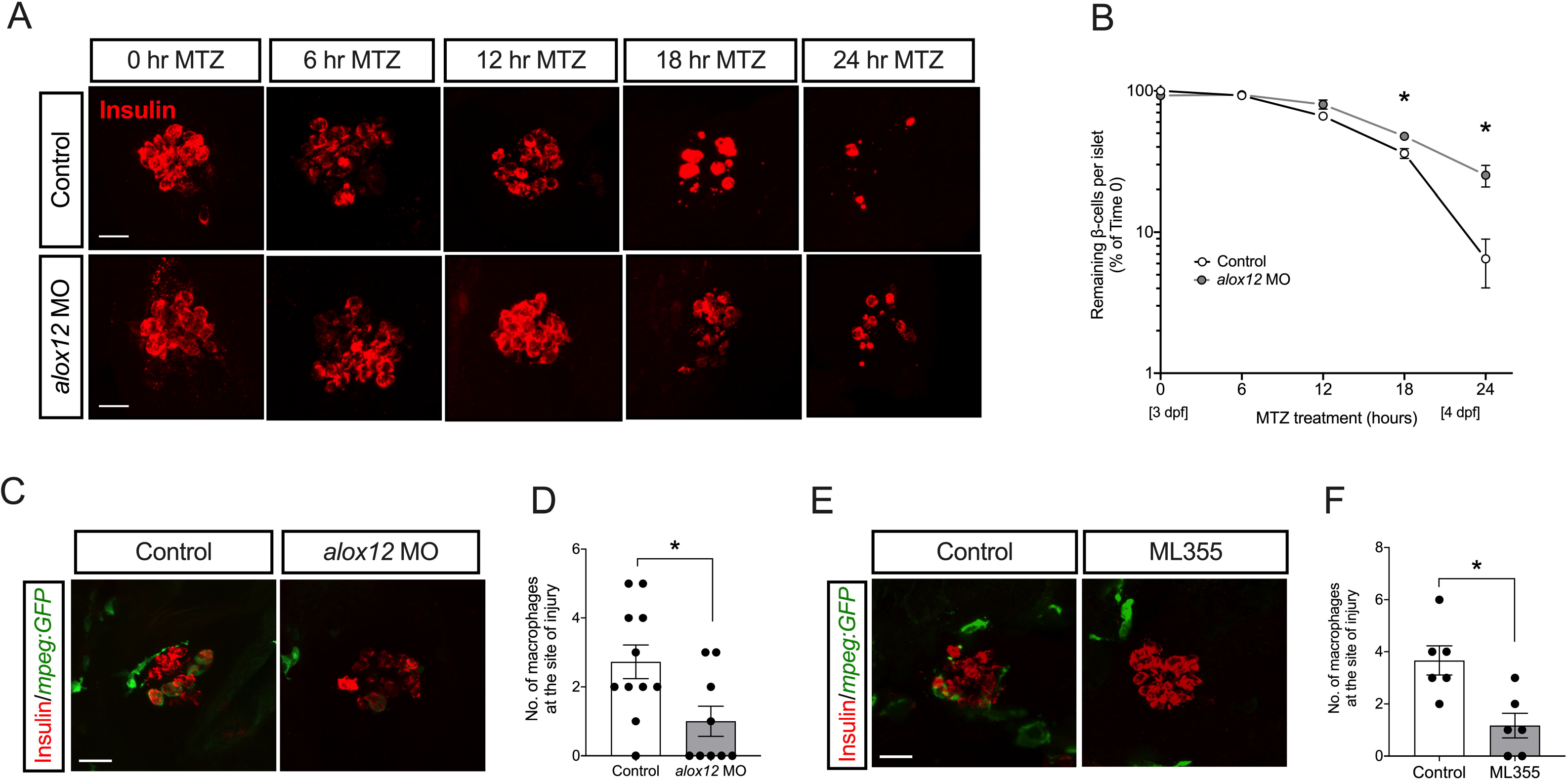
Depletion or inhibition of 12-LOX protects against β-cell loss in zebrafish. Zebrafish were treated with *alox12* morpholino (*MO*) or the 12-LOX inhibitor ML355 prior to treatment with MTZ at 3 dpf. (***A***) Representative images of islets from zebrafish stained for insulin (*β cells, red*); (***B***) Quantification of β-cell number in pancreatic islets of fish (expressed as % of β cells remaining relative to time 0) following MTZ treatment. N=6-8 fish per timepoint. *\*P*<0.05 for the timepoints indicated compared to control-treated fish; (***C***) Representative images of islets from control and *alox12* MO zebrafish stained for insulin (*β cells, red*) and GFP (*macrophages, green*) at 24 hours following MTZ treatment; (***D***) Quantification of the number of macrophages located at the site of injured islets from the experiment shown in *panel **C**. (**E***) Representative images of islets from control and ML355-treated zebrafish stained for insulin (*β cells, red*) and GFP (*macrophages, green); (**F***) Quantification of the number of macrophages located at the site of injured islets from the experiment shown in *panel **E***. Scale bar indicates 10 μm. In all panels, data are presented as mean□±□SEM. \**P* <0.05 for the comparisons shown.

To support our findings with the *alox12* MO, we next repeated these studies utilizing ML355, a small molecule inhibitor of 12-LOX [26]. As shown in Fig. 2E-F, when 12-LOX activity was inhibited by treatment of fish with 10 μM ML355, macrophage infiltration into the injured islets was reduced 3.1-fold relative to vehicle-treated controls. Collectively, the data in Fig. 2 suggest that 12-LOX is required for macrophage-directed β-cell damage in zebrafish, and raise the possibility for a similar role in β-cell damage in T1D.

### 12-LOX in macrophages is required for T1D progression in the mouse

To assess the applicability of our zebrafish findings to an established mammalian model of T1D, we next studied the non-obese diabetic (NOD) mouse model. We tested the hypothesis that 12-LOX in macrophages is responsible for the progression to T1D in NOD mice by generating a myeloid lineage-specific deletion of *Alox15* on the NOD background (*NOD:Alox15*^Δ*myel*^). NOD mice harboring the *Lyz2-Cre* allele were crossed to NOD mice harboring Cre recombinase sites (Loxp) flanking exons 2-5 of the *Alox15* gene [27,28]. To test for tissue specificity of the knockout, *Alox15* mRNA was measured in peritoneal cells (containing mostly macrophages), spleen, and islets using quantitative PCR. *NOD-Alox15*^Δ*myel*^ mice exhibited a significant reduction in *Alox15* expression in peritoneal cells compared to control littermates, whereas *Alox15* expression in spleen and pancreatic islets were unchanged (Fig. 3A).

**Figure 3:**
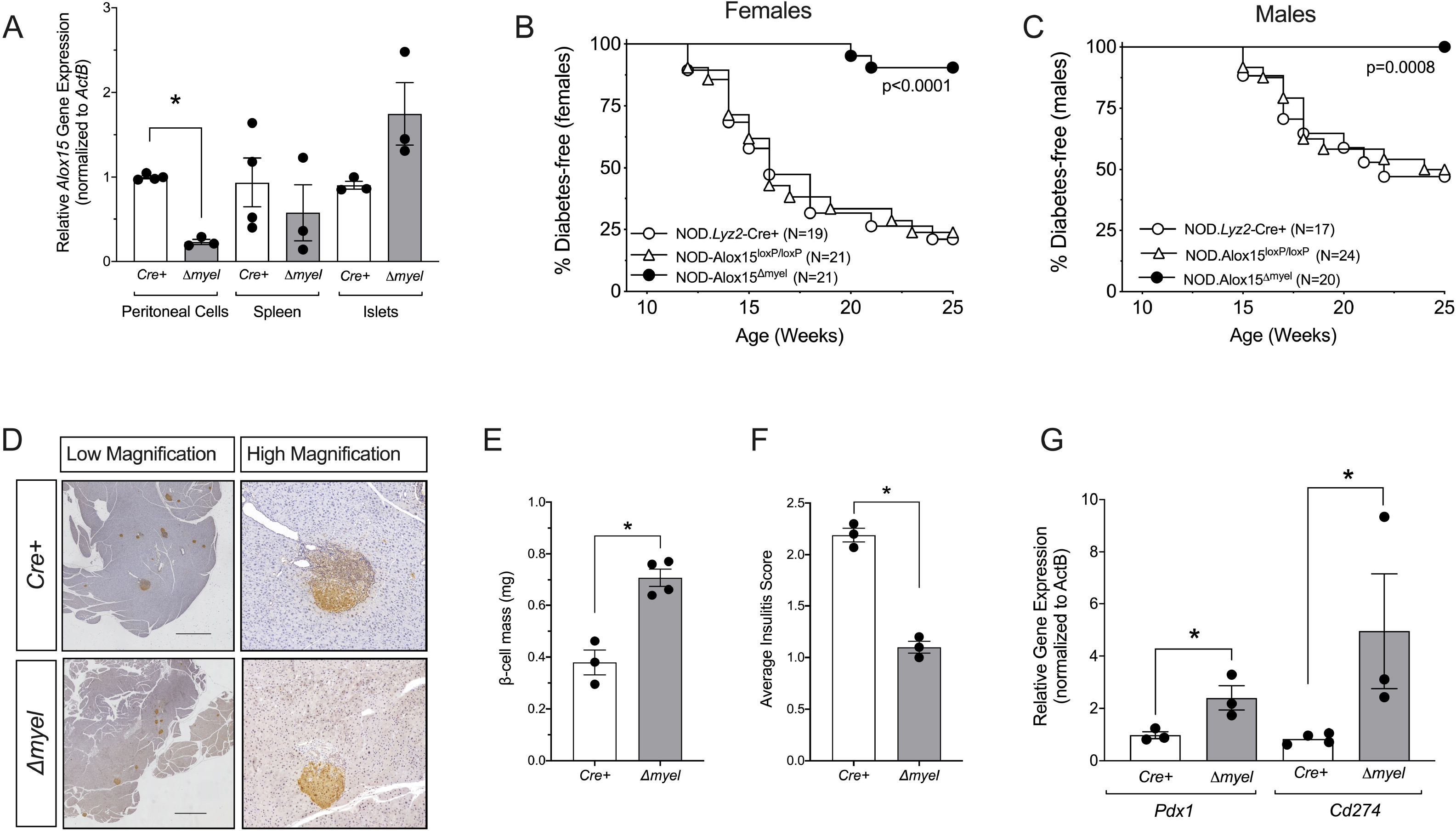
Protection from diabetes following myeloid-specific deletion of *Alox15* in NOD mice. NOD mice harboring the *Lyz2-Cre* allele were crossed to NOD mice harboring Cre recombinase sites (Loxp) flanking exons 2-5 of the *Alox15* gene to generate *NOD:Alox15*^Δ*myel*^ mice (Δ*myel*). *NOD:Lyz2-Cre* (Cre+) and *NOD:Alox15^Loxp/Loxp^* littermates were used as controls. (***A***) *Alox15* mRNA expression in peritoneal cells, spleen and isolated islets; (***B***) Diabetes incidence in female mice. Number (N) of mice per group is indicated. *P* value indicates significance by log-rank test; (***C***) Diabetes incidence in male mice. Number (N) of mice per group is indicated. *P* value indicates significance by log-rank test; (***D***) Representative immunohistochemical images of mouse pancreata from *Cre*+ and Δ*myel* mice immunostained for insulin (*brown*) and counterstained with hematoxylin (*blue*). Scale bar indicates 1000□μm; (***E***) β-cell mass in *Cre*+ and Δ*myel* mice; (***F***) Insulitis scoring from pancreata of *Cre*+ and Δ*myel* mice; (***G***) Gene expression in isolated islets from *Cre*+ and Δ*myel* mice. All data are presented as mean□±□SEM. \**P*<0.05 for the comparisons shown.

Next, we followed *NOD:Alox15*^Δ*myel*^ mice and control littermates (*NOD:Lyz2-Cre* and *NOD:Alox15^Loxp/Loxp^*) for the spontaneous development of T1D, defined as two consecutive morning blood glucoses >250 mg/dl. As shown in Fig. 3B, whereas 75-80% of female control littermates developed diabetes by 25 weeks of age, only 12.5% of female *NOD:Alox15*^Δ*myel*^ mice developed diabetes. Similar striking findings were observed in male mice: 50% of controls developed diabetes by 25 weeks of age, whereas 0% of *NOD:Alox15*^Δ*myel*^ developed diabetes (Fig. 3C). These findings suggest that the T1D-protective phenotype previously described in global *Alox15*-/- mice [18] might be ascribed at least in part to its effect in macrophages.

To characterize islet pathology in NOD mice, we next performed immunostaining of pancreas sections from *NOD:Alox15*^Δ*myel*^ mice and *NOD:Lyz2-Cre* controls at 8 weeks of age, a timepoint when insulitis is established but prior to the development of T1D. As evident in the representative immunohistochemical pancreas images in Fig. 3D, β-cell mass was significantly increased (quantitated in Fig. 3E) and the severity of insulitis was significantly reduced (quantitated in Fig. 3F) in *NOD:Alox15*^Δ*myel*^ animals compared to *NOD:Lyz2-Cre* controls (Fig. 3F). Isolated islets from *NOD.Alox15*^Δ*myel*^ mice revealed an increase in mRNAs levels of *Pdx1*, which encodes a key transcription factor that promotes β-cell function [29], and *Cd274*, which encodes PD-L1, an immune checkpoint protein that promotes suppression of immune responses in T1D [30] (Fig. 3G).

### Myeloid-specific loss of 12-LOX alters the macrophage and dendritic cell populations in pancreatic lymph nodes

Pancreatic lymph nodes are a site of local antigen presentation and their immune cell composition is reflective of the nature of prevailing autoimmunity [8]. To assess quantitatively if the autoimmune response was altered in *NOD:Alox15*^Δ*myel*^ mice, we collected pancreatic lymph nodes and performed flow cytometry for immune cell populations. Fig. 4A-C shows that populations of proinflammatory macrophages (F4/80+/TNF-α+) and myeloid-derived antigenpresenting dendritic cells (CD11c+/TNF-α+) were significantly reduced in total number and as a percentage of total cells in pancreatic lymph nodes of *NOD:Alox15*^Δ*myel*^ compared to *NOD:Lyz2-Cre* controls. Additionally, there were reductions in both macrophages and dendritic cells expressing the proinflammatory cytokine IL-1β (Supplementary Fig. S2A-B). These alterations in macrophage and dendritic cell populations were specific to the pancreatic lymph node, since these cell populations were unchanged in the spleen (Supplementary Fig. S3). Because lymph nodes are a site of antigen presentation by macrophages to CD4+ T cells, we also quantitated CD4+ cells in the pancreatic lymph nodes. As shown in Fig. 4C, we observed reductions both in total number of CD4+ cells and in CD4+ cells as a percentage of total cells in the pancreatic lymph nodes in *NOD:Alox15*^Δ*myel*^ compared to *NOD:Lyz2-Cre* controls.

**Figure 4:**
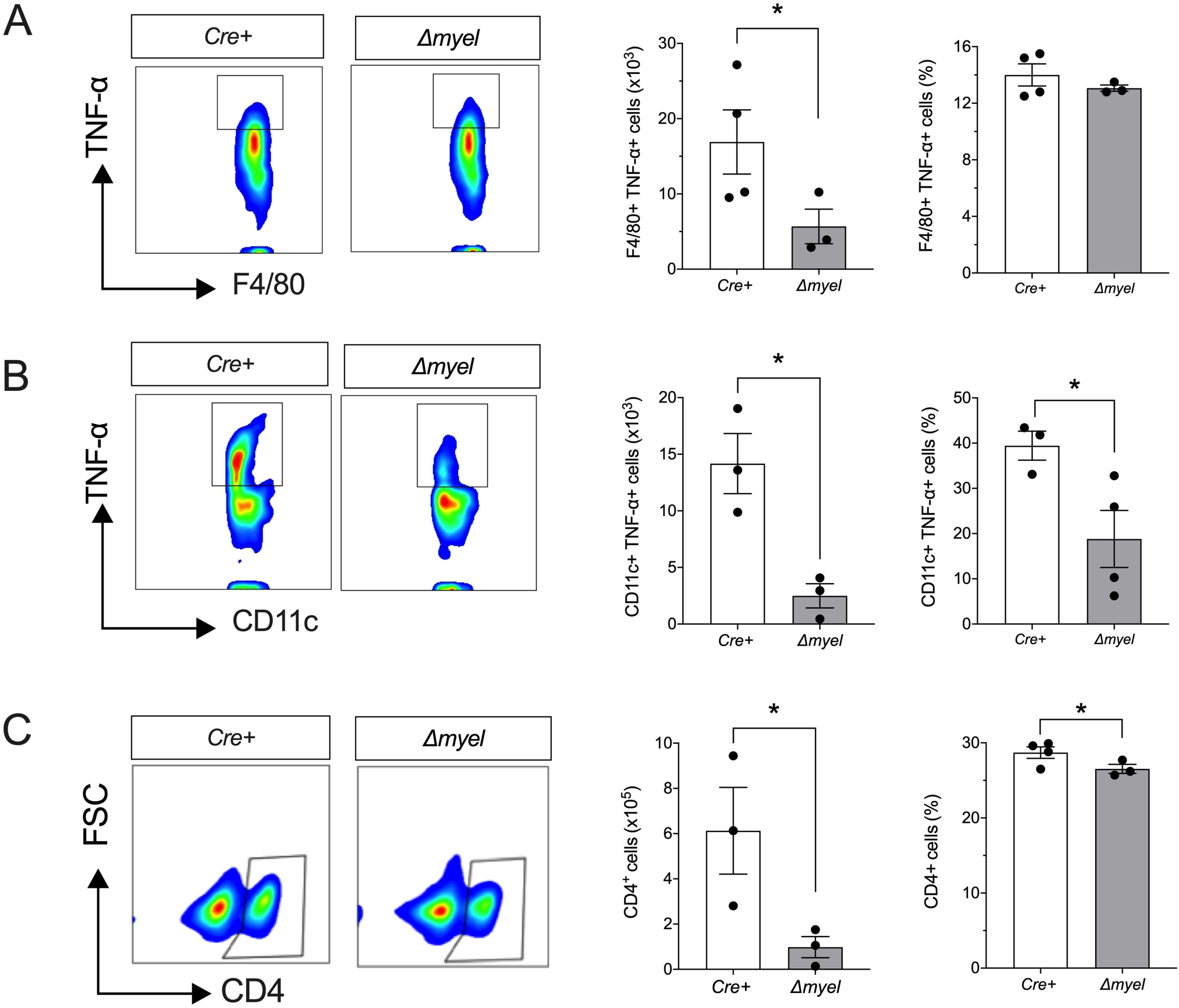
Reduced proinflammatory myeloid cell populations and CD4 T cells in pancreatic lymph nodes of *NOD:Alox15^Δmyel^* mice. Pancreatic lymph nodes were isolated from *NOD:Lyz2-Cre* (*Cre*+) control mice and *NOD:Alox15*^Δ*myel*^ (Δ*myel*) mice at 8 weeks of age and subjected to flow cytometry analysis. (***A***) Representative contour plot showing gating of F4/80+ TNFα+ cells (*left*), total number of F4/80+ TNFα+ cells (*middle*), and F4/80+ TNFα+ cells as a percentage of total cells (*right*); **(*B*)** Representative contour plot showing gating of CD11c+ TNFα+ cells (*left*), total number of CD11c+ TNFα+ cells (*middle*) and CD11c+ TNFα+ cells as a percentage of total cells (*right*); (***C***) Representative contour plot showing gating of CD4+ cells (*left*), total number of CD4+ cells (*middle*), and CD4+ cells as a percentage of total cells (*right*). All data are presented as mean□±□SEM. \**P* <0.05 for the comparisons shown.

### 12-LOX promotes macrophage migration

To clarify the mechanisms underlying the less aggressive macrophages in the absence of 12-LOX, we next asked if 12-LOX governs the polarization of macrophages to a proinflammatory state. We isolated peritoneal macrophages from *Alox15*-/- mice or their WT littermates, performed polarization studies in vitro, then monitored phenotypes by flow cytometry. Macrophages were polarized to the classical proinflammatory “M1-like” state using a combination of lipopolysaccharide (LPS) and IFN-γ or to the alternative anti-inflammatory “M2-like” state using IL-4 [31]. The macrophages from both WT and *Alox15*-/- mice showed indistinguishable propensity to polarize to an M1 state, as assessed by flow cytometry of M1 marker iNOS (Supplementary Fig. S4A), by IL-6 secretion (Supplementary Fig. S4B), and by mRNA expression of *Nos2, Il6* and *Il12* (Supplementary Fig. S4C). Similarly, upon alternative polarization, macrophages from both WT and *Alox15*-/- mice showed no differences by flow cytometry in the M2 marker CD206 (Supplementary Fig. S5A), by IL-10 secretion (Supplementary Fig. S5B), or by mRNA expression of *Arg1, Il10* and *Tgfb* (Supplementary Fig. S5C). Taken together, these data indicate that 12-LOX activity is not required for cytokine-induced polarization of macrophages.

Next, we addressed if 12-LOX is required for the ability of cells of the myeloid lineage to migrate. Migration is a critical factor in the ability of myeloid cells to carry out immune surveillance, locate to sites of injury, and present antigen. To address the potential role of 12-LOX in macrophage migration, we first leveraged the tailfin injury model in *Tg(mpeg:eGFP)* transgenic zebrafish. *alox12* MO and control fish underwent tailfin injury at 3 dpf, and the number of macrophages migrating to the site of injury was quantitated. In control fish, we observed the expected migration of macrophages to the site of injury within 6 hours, whereas in *alox12* MO fish there was a significant 38% reduction in the number of macrophages at the site of injury (Fig. 5A). As an alternate approach, and to confirm that this finding requires the catalytic activity of 12-LOX, we performed a similar experiment using the well-characterized small molecule 12-LOX inhibitor ML355 [26]. For this experiment, we pre-treated fish for 2 hours with either vehicle or 10 μM ML355, and then performed tailfin injury. Similar to what we observed with the morpholino knockdowns, ML355 treatment significantly decreased macrophage migration towards injured sites by 37% as compared to vehicle-treated control embryos (Fig. 5B).

**Figure 5:**
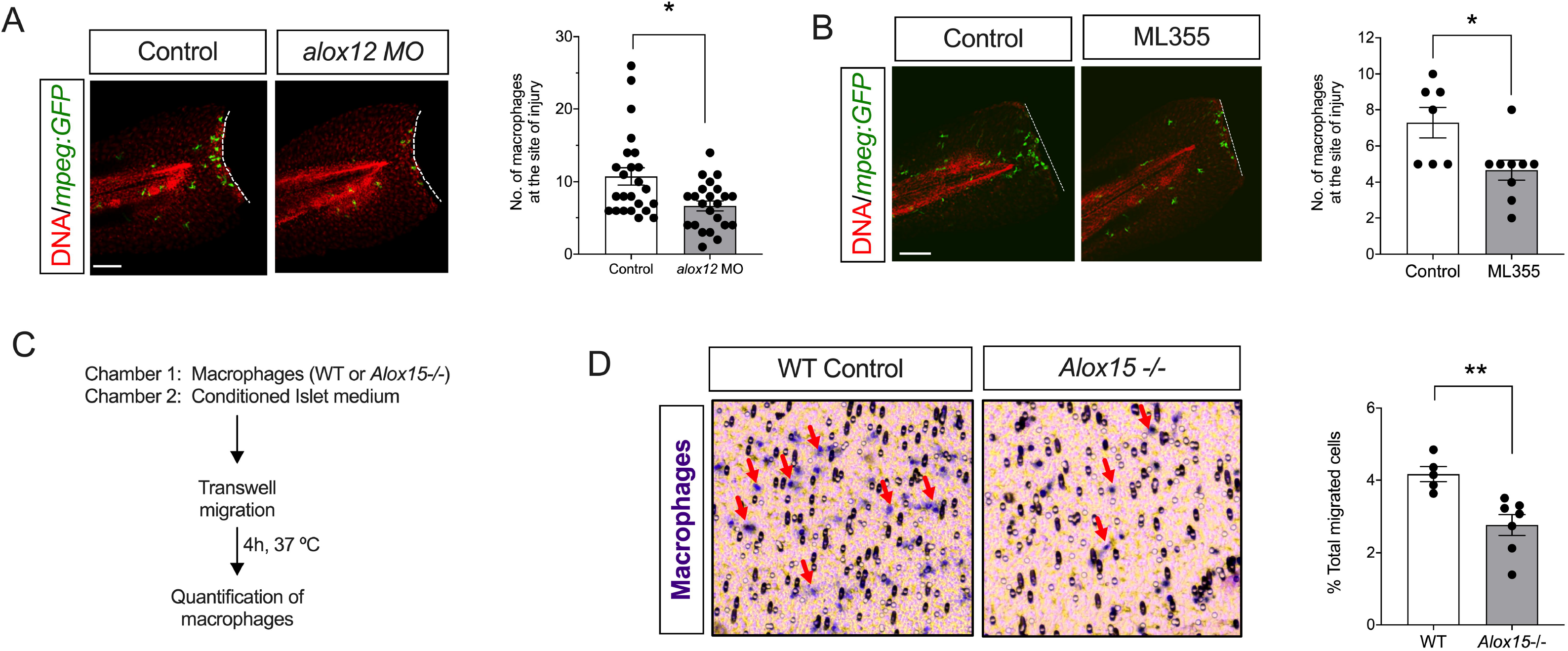
Depletion or inhibition of 12-LOX impairs macrophage migration in zebrafish and mice. (***A***) Control and *alox12* MO-treated *Tg(mpeg:GFP)* zebrafish underwent tailfin injury at 3 dpf. Representative images of injured zebrafish tails stained with GFP (*macrophages, green*) and TO-PRO3 (*nuclei, red*) are shown on the *left* and quantitation of migrating macrophages is shown on the *right*; (***B***) *Tg(mpeg:GFP)* zebrafish were treated with vehicle or 10 μM ML355 and then underwent tailfin injury at 3 dpf. Representative images of injured zebrafish tails stained with GFP (*macrophages, green*) and TO-PRO3 (*nuclei, red*) are shown on the *left* and quantitation of migrating macrophages is shown on the *right*; (***C***) Description of the chemotaxis assay in vitro using wild-type (WT) and *Alox15*-/- mouse peritoneal macrophages; (***D***) Representative images of the porous membrane showing migrating methylene blue-stained macrophages (*red arrows*) is shown on the *left*, and quantitation of the number of migrating macrophages is shown on the *right*. Scale bar indicates 50 μm. All data are presented as mean□±□SEM. \**P* <0.05 for the comparisons shown.

To verify that our findings were specific to 12-LOX in macrophages and to confirm their applicability to mammals, we isolated peritoneal macrophages from both *Alox15*-/- mice and their littermate controls and performed migration assays *in vitro* using transwell chambers. To mimic conditions seen in T1D, we added conditioned culture media from proinflammatory cytokine-treated (IL-1β, IFN-γ, TNF-α) mouse islets to induce migration (Fig. 5C). As shown and quantitated in Fig. 5D, we observed a significant reduction in the number of *Alox15*-/- peritoneal macrophages that transited the transwell membrane compared to WT control macrophages. These results, consistent with the zebrafish studies, indicate that 12-LOX in macrophages contributes to their ability to migrate under tissue damage/inflammatory conditions.

### Chemokine receptor CXCR3 lies downstream of 12-LOX activity

Prior studies suggest that 12-LOX and its product 12-HETE alters chemokine receptor expression [32–34]. We hypothesized that the macrophage migratory defect we observed in the absence of 12-LOX may be attributable to loss of one or more of these chemokine receptors. We measured the mRNA expression of chemokine receptors *Cxcr1, Ccr2*, and *Cxcr3* implicated in macrophage migration in peritoneal macrophages from *Alox15*-/- and control littermate mice. Of the receptor-encoding mRNAs examined, only *Cxcr3* levels were significantly reduced in *Alox15*-/- macrophages (Fig. 6A). Flow cytometry of peritoneal macrophages confirmed that the cell surface expression of CXCR3 protein is significantly reduced in *Alox15*-/- macrophages compared to WT controls (Fig. 6B).

**Figure 6:**
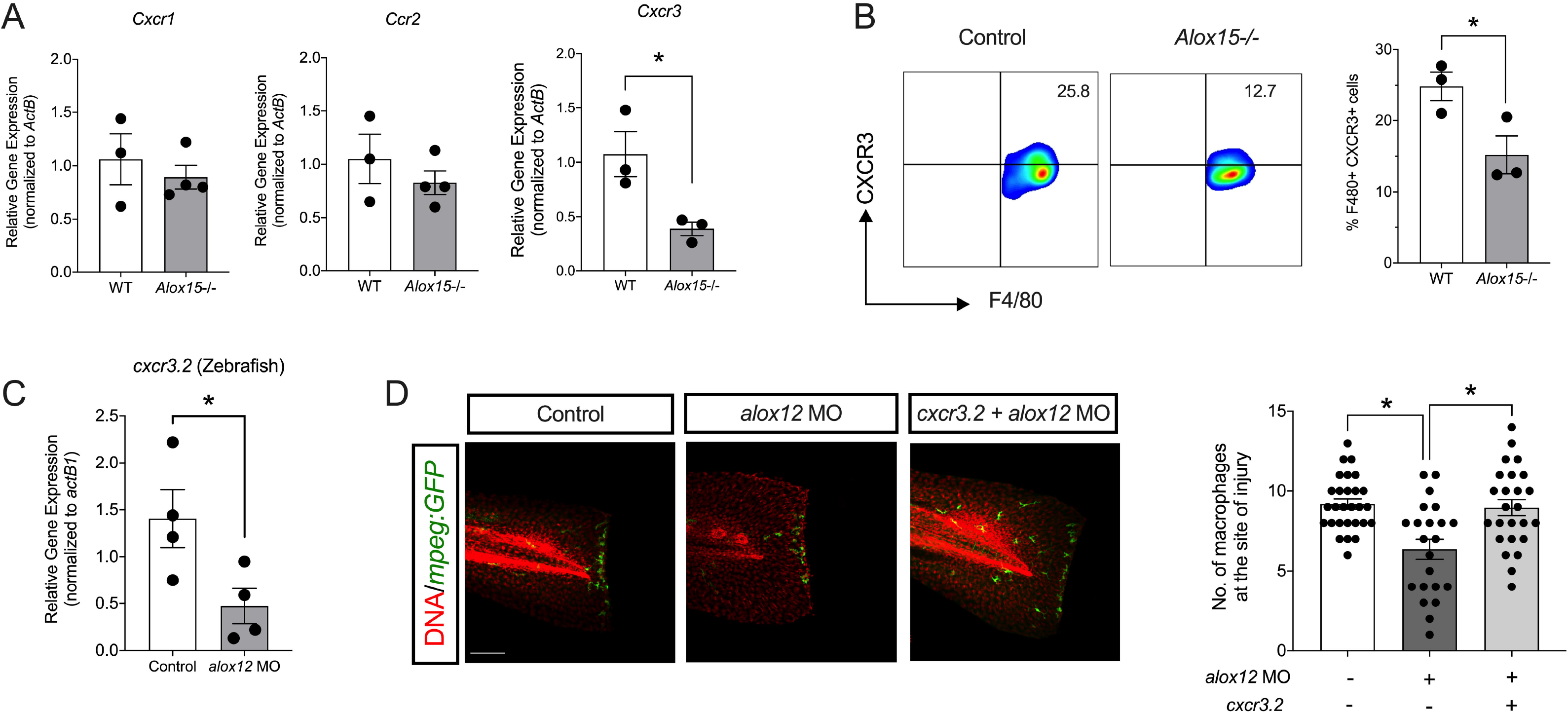
CXCR3 is reduced in the absence of 12-LOX. (***A***) Chemokine receptor mRNA expression in macrophages from wild-type (WT) and *Alox15*-/- mice; (***B***) Representative contour plots showing gating of F4/80+ CXCR3+ cells are shown on the *left* and F4/80+ CXCR3+ cells as a percentage of total peritoneal cells from WT and *Alox15*-/- mice is shown on the *right*; (***C***) *cxcr3.2* mRNA expression from whole lysates of control- and *alox12* MO-treated zebrafish; (***D***) Representative images of injured *Tg(mpeg:GFP)* zebrafish tailfins stained with GFP (*macrophages, green*) and TO-PRO3 (*nuclei, red*) are shown on the *left*, and corresponding quantitation of macrophages at the tail injury site in the control, *alox12* MO injected fish and the fish co-injected with *alox12* MO and vector for macrophage specific overexpression of *cxcr3.2* is shown on the *right*. Scale bar indicates 100 μm. Data are presented as mean□±□SEM. \**P* <0.05 for the comparisons shown.

To interrogate if the loss of CXCR3 accounts, at least in part, for the defect in macrophage migration observed in the absence of 12-LOX, we returned to zebrafish. As shown in Fig. 6C, 12-LOX depletion in *alox12* MO zebrafish resulted in a similar loss of expression of the zebrafish ortholog *cxcr3.2*. Next, we transgenically expressed a *cxcr3.2* coding sequence specifically in zebrafish macrophages (under control of the *mpeg* promoter) to determine if reexpression of this gene was sufficient to rescue the migratory defect in *alox12* MO fish. As shown and quantitated in Fig. 6D, *alox12* MO fish injected with a non-coding control vector exhibited the expected reduction in macrophage migration in the tailfin injury assay compared to controls; however, *alox12* MO fish injected with the *cxcr3.2*-containing vector showed complete rescue of macrophage migration. These results suggest that the reduction of *cxcr3.2* in 12-LOX-depleted zebrafish likely accounts for the defect in macrophage migration.

## DISCUSSION

In this study, we leveraged the power of a lower model organism and applied these observations to an established mammalian model of T1D to interrogate the participation of myeloid-derived cells in the pathogenesis of insulitis and diabetes progression. Cells of myeloid lineage give rise to a host of circulating cell types, most notably cells of the innate immune system that include monocytes, macrophages, and antigen-presenting dendritic cells. Although such cells have been implicated in the pathogenesis of T1D, and prior studies have attempted myeloid-specific deletions in T1D mouse models [35], we believe our findings are the first to directly demonstrate how manipulation of a signaling pathway specifically in myeloid cells can alter T1D pathophysiology. Key findings from our study suggest (a) that macrophages are active participants in β-cell dysfunction and loss in models of both islet injury and autoimmunity, (b) that 12-lipoxygenase establishes a pro-diabetogenic phenotype of macrophages and dendritic cells that subsequently affects the nature of insulitis and eventual susceptibility to T1D, and (c) that the *Tg(ins:NTR)* zebrafish transgenic line provides a highly genetically manipulable platform wherein crosstalk between macrophages and β cells can be modeled and vetted.

Islet-resident myeloid cells have been implicated in the pathogeneses of both T1D and T2D [14,36]. In NOD mice, prior studies have emphasized the role of such resident cells as initiators of the autoimmune response, owing perhaps to their role as antigen presenting cells. These studies focused on depletion of macrophages, using either clodronate liposomes [11,13,15,16] or neutralization of a receptor (CSF1 receptor) promoting their development [14]. Studies of Unanue and colleagues [14] demonstrated that delivery of a neutralizing antibody against the colony stimulating factor 1 (CSF1) receptor at 3 weeks of age in NOD mice resulted in the depletion of islet-resident macrophages and was accompanied by reduction in CD4 T cells and dendritic cells in the insulitic milieu and protected against diabetes. These results align with ours, in which deletion of *Alox15* in macrophages on the NOD background resulted in significant reduction in insulitis, CD4 T cells, and protection from diabetes in both male and female mice. Moreover, our findings in vivo that myeloid-specific depletion of *Alox15* reduces pro-inflammatory dendritic cells and enhances expression of the mRNA encoding the immune checkpoint protein PD-L1 suggests that the effects we observed may be related to altered antigen presentation and or adaptive immune cell activation, respectively. Collectively, our results emphasize that early intervention in the activity of myeloid cells imparts a disease-modifying effect in T1D, but more importantly that depletion of the islet-resident populations of these cells is not required to achieve this effect.

An important new finding in our studies is the identification that 12-LOX controls a signaling pathway that affects chemokine receptor expression in macrophages. The gene encoding 12-LOX (*Alox15*) is expressed in islet β cells and macrophages, but not T cells or B cells of the adaptive immune system [37]. Although *Alox15* deletion was previously shown to protect against T1D in NOD mice [18], it has remained unclear if the effect could be attributed to its expression islets, myeloid cells, or both, particularly since proinflammatory cytokines and their signaling are affected by loss of *Alox15* in both cell types ([28,37]. Our studies in zebrafish provided initial evidence for a defect in macrophage function upon loss of the zebrafish ortholog, since macrophage entry into the islet and subsequent engulfment of β cells appeared defective in *alox12* morphants upon MTZ injury. Though it is tempting to speculate that the phenotype in these morphants was attributable to the loss of *alox12* in macrophages, our studies of conditional *Alox15* deletion in myeloid-derived cells in NOD mice proved more conclusive in this regard. Importantly, whereas prior studies of *Cre* recombinase expression on the NOD background leave some doubt as to the effect of the deletion vs. the misexpression of *Cre* recombinase on diabetes outcome [38], we show rigorously here that inclusion of littermate controls containing either the homozygous *Loxp* alleles or expressing *Cre* recombinase develop T1D at the expected frequencies in both sexes. Therefore, we believe that the remarkable protective phenotype of NOD mice harboring myeloid cell-specific loss of the *Alox15* alleles is attributable directly to loss of 12-LOX activity in these cells.

The mechanism by which 12-LOX promotes T1D progression has been variably attributed to cytokine signaling, oxidative stress, and cellular apoptosis induced by its major arachidonic acid-derived eicosanoid 12-hydroxyeicosatetraenoic acid (12-HETE) [17]. The majority of these prior studies were performed in islets and β cell-derived cell lines. Here, we demonstrate new mechanisms by which 12-LOX impacts myeloid cell function in T1D. We show that 12-LOX is not required for the apparent polarization of macrophages to pro- or antiinflammatory states, suggesting that its effects may be more specific to functional duties commonly ascribed to myeloid cells. The chemokine receptor CXCR3 and its ligands CXCL9, CXCL10, and CXCL11 serve as part of the chemoattractant response during inflammation and immunity. Here and in prior studies, CXCR3 has been shown to be expressed on myeloid-derived cells [39], and may be required for local migration for immune surveillance, antigen presentation, and tissue damage clearance. Prior studies on the role of CXCR3 in the context of NOD mice have shown conflicting results, with some studies suggesting that its deficiency protects against diabetes [40] and others suggesting its deficiency accelerates diabetes [41]. A complexity in these prior studies is the global nature of deletion and the differences in diabetes induction in these animal models. Although our studies do not entirely resolve this issue, they do indicate a critical role of this receptor for a fundamental function of myeloid cells. We show that 12-LOX is required for cellular migration, likely through the expression of CXCR3 in mice and its ortholog CXCR3.2 in zebrafish, where re-expression of *cxcr3.2* specifically in macrophages restored the migratory capacity. That the migratory defect can be rescued by transgenic re-expression emphasizes that the effect of 12-LOX is exerted at the transcriptional, rather than posttranscriptional, level. This finding is especially relevant as 12-LOX is an enzyme with no known transcriptional function *per se*. It remains possible that this transcriptional effect is the result of the downstream activity of GPR31, a receptor for 12-HETE [25,42]. Further studies of *Gpr31* mutant zebrafish and mice will be required to address this issue.

A final major implication of our study is the relevance of zebrafish. We present a model system in which macrophages play a central role in destruction of β cells in a transgenic *Tg(ins:NTR)* line of zebrafish. In the absence of adaptive immune cell involvement in our zebrafish model, it would be inaccurate to claim that this system models T1D. Nevertheless, this model system exhibits some features that allow its use as a platform for the study the dynamics between macrophages and β cells. We showed that depletion of macrophages in zebrafish using clodronate preserved pre-existing β cells following MTZ treatment of zebrafish, yet prevented the formation of neogenic β cells—the combined effect of which may have resulted in only the modest effect on glycemia that we observed. On the one hand, these findings reflect similar studies in NOD mice in which clodronate treatment preserved β cells and glycemia [11,11,16] and, on the other, they support studies in mice that macrophages promote β-cell proliferation tissue regeneration [43,44]. This seemingly dichotomous finding likely reflects the differential involvement of pro-inflammatory (“M1”) and anti-inflammatory (“M2”) macrophages, which are not differentiated by clodronate treatment. By contrast, our studies with *alox12* MO and the 12-LOX inhibitor ML355 appears to target the “pro-diabetogenic” (presumably M1-like) macrophages, findings that are supported by our tissue-specific knockout studies in NOD mice. In ongoing studies, our laboratory is interrogating the potential existence of different macrophage phenotypes in zebrafish, the implications of which would broaden further the applicability of zebrafish models for interrogating innate immune-mediated tissue injury and repair.

In conclusion, our findings support a role for macrophages and dendritic cells in the initiation of T1D, but more importantly implicate a central role for 12-LOX in promoting the initial innate immune response during diabetes pathogenesis. Fig. 7 shows a schematic representation of the findings of our study in the context of the dendritic cell-β cell interactions that govern the pathophysiology of T1D. We propose that 12-LOX in pro-inflammatory macrophages and dendritic cells promotes the expression of *Cxcr3* (possibly via GPR31) to permit cellular migration, antigen acquisition and presentation, as well as production of proinflammatory cytokines (IL-1β, TNF-α). The deletion or inhibition of 12-LOX in the early phases of T1D impairs these processes, thereby reducing antigen acquisition and presentation, insulitis, and β-cell loss. This model does not explicitly exclude an independent role for 12-LOX in β cells, where it promotes pro-inflammatory signaling leading to cellular stress and apoptosis. Whether deletion of *Alox15* in β cells might independently disrupt this interaction between cells remains an ongoing goal of our laboratory. Some limitations of our study are worth noting. Our study deleted 12-LOX from the inception of *Lyz2* expression in all myeloid cells during mouse development; hence, it remains unclear (a) if its role beyond macrophages and dendritic cells (such as neutrophils) might also contribute to the phenotype observed, and (b) if the loss of *Alox15* during myeloid cell development might have inherently prevented the acquisition of a pro-diabetogenic phenotype before even the initiation of the disease process. The latter point could have implications for the timing of targeted therapeutics. Ongoing studies in the lab are focused on the timing of *Alox15* deletion relative to disease onset and the translation of these studies to human disease and humans.

**Figure 7:**
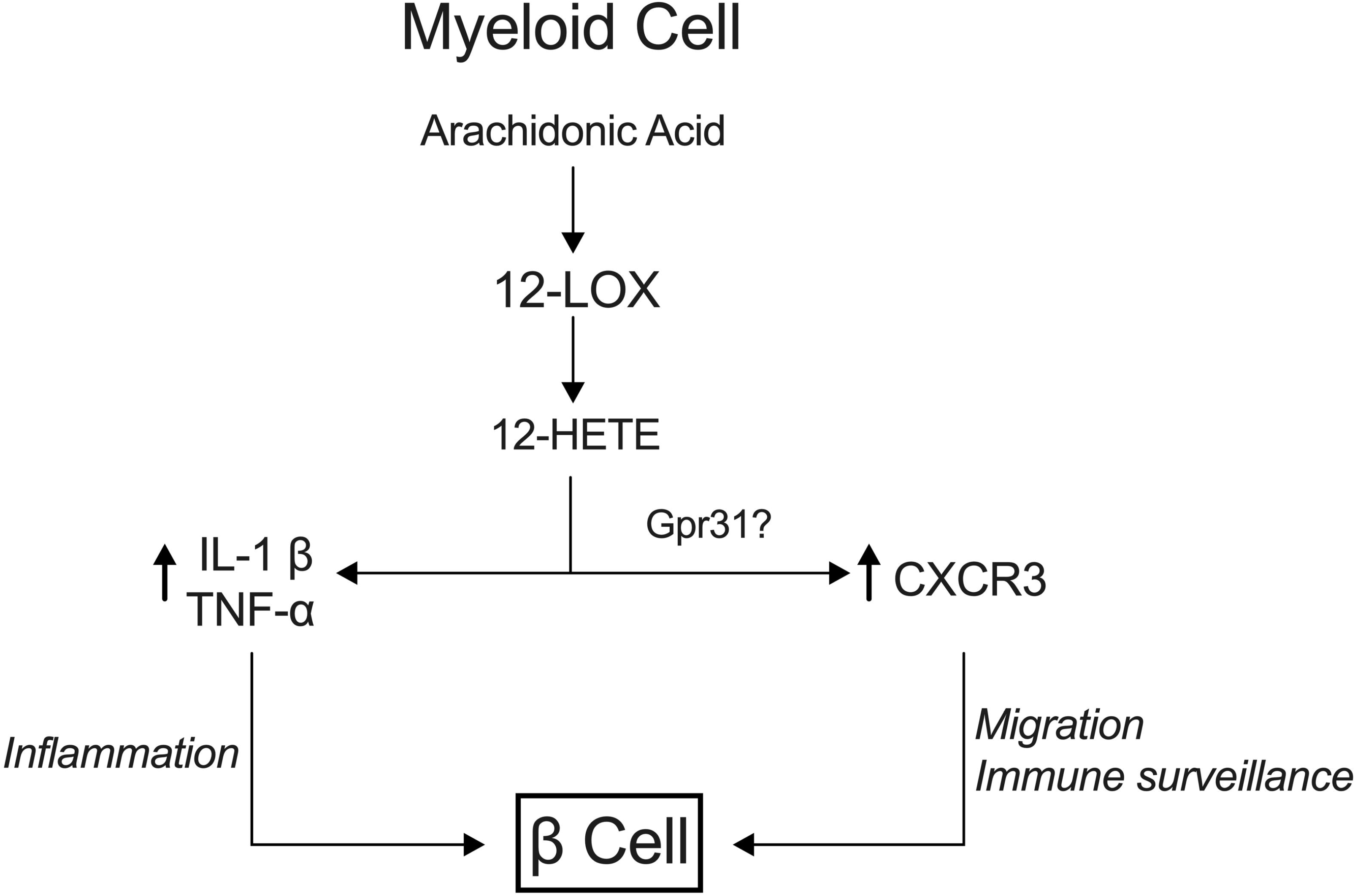
Model for 12-LOX regulation of macrophage inflammation, migration and surveillance. The model shown depicts the proposed role of 12-LOX in innate immune myeloid cells during the pathogenesis of type 1 diabetes. The activity of 12-LOX catalyzes the production of 12-hydroxyeicosatetraenoic acid (12-HETE) from membrane phospholipid-derived arachidonic acid. Elevations in 12-HETE levels result in increased production of proinflammatory cytokines (e.g. IL-1β, TNF-α) and enhanced expression of CXCR3, which lead, respectively, to islet β-cell inflammation and enhanced migration and immune surveillance.

## METHODS

### Zebrafish and mouse strains and maintenance

Zebrafish (*D. rerio*) were maintained at 28.5 °C in a recirculating aquaculture system enclosed in a cabinet and subjected to a 14 h:10 h light-dark cycle according to institutional protocols approved by both the Indiana University and University of Chicago Institutional Animal Care and Use Committees. The following transgenic lines were used in the experiments: *Tg(ins:NTR)^s950^* [45], *Tg(mpeg1:eGFP)^gl22^* [21], and *Tg(ins:Kaede)^s949^* [46]. Heterozygous outcrossed embryos bearing relevant transgenic alleles were collected at spawning and maintained in a 28.5°C incubator in petri dishes with buffered egg water [0.1% instant ocean salt, 0.0075% calcium sulfate supplemented with 0.003%1-Phenyl-2-thiourea (PTU; Acros) to prevent pigmentation in all embryos. At 3 days post fertilization (larval stage), the transgenic zebrafish were genotyped by epifluorescence using a Leica M205FA dissecting microscope. The *alox12* morpholino (MO) targeting the translational start was described previously [25]. A *cxcr3.2*-containing expression vector was generated by inserting the coding sequence of *D. rerio cxcr3.2* gene under control of the *D. rerio* macrophage-specific *mpeg1* promoter (*mpeg1:cxcr3.2*).

All mouse experiments were performed under pathogen-free conditions through protocols approved by the Indiana University, University of Chicago, and Eastern Virginia Medical School Institutional Animal Care and Use Committees. *Alox15*+/- mice on the *C57BL/6J* background were purchased from Jackson Laboratories and maintained and bred inhouse. For the experimental controls, we utilized wild-type (WT) littermates from breedings. *C57BL/6J.Alox15^loxP/loxP^* [27,28] and *C57BL/6J.129P2-Lyz2^tm1(cre)lfo^/J (Lyz2-Cre*) were backcrossed onto the *NOD.ShiLt/J* background using speed congenics services provided through Jackson Laboratories. After successful backcrossing, *NOD.Alox15^loxP/loxP^* mice were crossed with *NOD.Lyz2-Cre* mice to generate breeding colonies. All controls for NOD mouse experiments were littermates (either *NOD.Alox15^loxP/loxP^* or *NOD.Lyz2-Cre*). Diabetes incidence was monitored as described previously [47].

### Zebrafish manipulations

For the β-cell injury assay, zebrafish were washed with egg water and then treated with 7.5 mM metronidazole (MTZ) (Sigma) prepared in egg water or egg water alone as described previously [22]. Following MTZ treatment, fish were washed with egg water and allowed to recover for the times indicated. For the tailfin injury assay, fish were transiently paralyzed with 0.01% tricaine (Sigma) in egg water to restrict their movements, and the distal tip of the tail fin was amputated with a scalpel. At the end of each experiment, fish were fixed, deyolked, and immunostained as previously described [48]. The following concentrations of primary antibodies were used: 1:200 guinea pig anti-insulin (Invitrogen), 1:200 chicken anti-GFP (Aves Labs). Primary antibodies were detected with 1:500 dilutions of complementary Alexa-conjugated secondary antibodies (Jackson ImmunoResearch). DNA was stained with 1:500 TO-PRO3 (Thermo Fisher). After staining, fish were mounted on charged glass slides in VECTASHIELD (Vector Labs) and confocal imaging was performed with an LSM700 or LSM800 microscope (Zeiss). For macrophage depletion studies, fish were first sedated with 0.01% tricaine, mounted in 2.5% methylcellulose (Electron Microscopy Sciences) and then injected trans-pericardially with 7-10 nL clodronate liposomes (Encapsula Nano Sciences) 24 hours prior to experimentation. For inhibitor studies, fish were pretreated with 12-LOX inhibitor (ML355) or vehicle (0.1% DMSO) for 2 hours.

For measurement of glucose levels, whole fish were homogenized in 500 μL glucose assay buffer provided in the Glucose Colorimetric Assay Kit (Biovision). Samples were then centrifuged at 2000 *xg* for 5 minutes at room temperature and the supernatant was used in duplicate for the glucose assay following manufacturer’s protocol. The colorimetric assay was measured using a SpectraMax iD5 multi-mode microplate reader (Molecular Devices) at 405 nm.

### Immunohistochemistry and β-cell mass

Pancreata from at least five different mice per group were fixed in 4% paraformaldehyde, paraffin embedded, and sectioned onto glass slides. Pancreata were immunostained using rabbit anti-insulin (1:1000; ProteinTech). β-cell mass was calculated as previously detailed [49]. Insulitis was scored as previously described [47] using the following scoring criteria: 1) no insulitis, 2) infiltrate < 50% circumference, 3) infiltrate > 50% circumference, 4) infiltration within islet.

### Primary cell isolations, incubations, and analyses

Islets from mice were isolated as previously described [50]. Mouse peritoneal macrophages were isolated as described [51] immediately after euthanasia by injecting ice-cold RPMI into the peritoneal cavity using a 25g needle. The injected RPMI was then removed. The isolated cells were lysed with RBC lysis buffer (eBioscience) to remove red blood cells. Naïve T cells were isolated and purified from spleen and lymph nodes of *C57BL/6J* mice using EasySep™ Mouse CD4+ T Cell Isolation Kit (Stemcell Technologies).

For polarization studies in vitro, isolated peritoneal macrophages stimulated with 10 ng/ml LPS and 25 ng/ml IFN-γ (for M1 polarization), 10 ng/mL IL4 (for M2 polarization), or media control for 16 hours. Peritoneal macrophages from NOD mice were treated with 100 ng/mL PMA (Sigma), 500 ng/ml Ionomycin (Sigma) and Golgi Stop Plug (1:1000; BD Pharmigen) prior to immunostaining. To stain for surface antigens, cells were incubated with antibodies to F4/80 (BM-8, Biolegend), CD11c (HL3, BD Pharmigen), CD4 (RM4-5, Biolegend), CD80 (16-10A1, Biolegend), CD86 (GL-1, Biolegend) and MHC-II (M5/114.15.2, Biolegend) or the appropriate isotype controls for 30 min. For cytokines staining, the cells isolated were stimulated with 100ng/mL PMA (Sigma), 500ng/ml Ionomycin (Sigma) and Golgi Stop Plug (1:1000; BD Pharmigen). Then cells were permeabilized using Cytofix/Cytoperm (BD Pharmigen) and incubated with antibodies for TNF-α (Biolegend), IL-1β (Thermo Fisher). IL-17 (TC11-18H10, BD Pharmigen), IFN-γ (XMG12, BD Pharmigen) and FoxP3 (MF23, BD Pharmigen). All antibodies were used 1:100 dilution. Cells were filtered and acquired on the AttuneTM NxT Flow Cytometer or a FACS Canto II cytometer (BD). Data were analyzed using FlowJo software (Tree Star). Supernatant from stimulated peritoneal macrophages was collected and IL-6 and IL-12 levels were measured by ELISA according to the manufacturer’s instructions (eBiosciences) and read on a SpectraMax iD5 multi-mode microplate reader (Molecular Devices).

### Gene expression analysis by quantitative PCR

RNA was isolated from zebrafish or from mouse tissue using the RNeasy Plus Micro Kit (Qiagen) and was used to prepare cDNA using a commercial kit (Applied Biosystems). Quantitative RT-PCR was run using the SsoFast EvaGreen Supermix kit (Biorad) on a Quantstudio 3 thermocycler (Applied Biosystems). The average Ct value of three replicates was calculated and normalized to β-actin.

#### Transwell chemotaxis assay

A 96-well chemotaxis system (ChemoTx, 8 μm filter pore size; Neuro Probe) was loaded with conditioned media from islets isolated from *C57BL/6J* mice. Isolated islets were treated with a pro-inflammatory cytokine cocktail (50 ng/mL TNF-α, 25 ng/mL IL-1β, 100 ng/mL IFN-γ) for 24 hours after isolation to generate conditioned media. The conditioned media was collected and loaded in the bottom chamber of the chemotaxis system. WT or *Alox15*-/- macrophages (0.5 × 10^5^ cells) were added to the upper chamber and migration to the lower chamber was measured after incubation for 4 hours at 37 °C. After incubation, non-migrated cells were washed while the filter side containing migrated cells were stained with Kwik-Diff Solution kit (Themo-Shandon). The filter was mounted on the slides, and the number of migrated macrophages was quantified by manual counting under an LSM800 microscope (Zeiss).

### Statistical Analysis

All data are presented as mean□±□standard error of the mean (SEM). The data analyses were performed using the GraphPad Prism 9 software package. Significant differences between the mean values were determined using Student’s t-test, where two means were compared, and one-way analysis of variance (ANOVA) followed by post hoc Tukey’s test when more than two means were compared. The differences were considered statistically significant at *p<0.05*.

## Acknowledgements

This research was supported by the National Institutes of Health grants R01 DK060581 (to RGM), R01 DK 105588 (to RGM and JLN), and F30 DK122681 (to SI), by a JDRF postdoctoral fellowship (to MHP), and a DeVault fellowship (to AK). Research core services were provided by National Institutes of Health grants P30 DK097512 (to Indiana University) and P30 DK020595 (to University of Chicago). The authors wish to acknowledge Dr. Marcia McDuffie (University of Virginia) for critically reading the manuscript and providing suggestions for improvements.

## Author Contributions

Conceptualization: AK, JLN, MAM, SAT, RGM, and RMA; Methodology: AK, ARP, MW, SI, MHP, KSO, LG, MAM, SAT, and RMA; Data analysis: AK, ARP, MW, JLN, MAM, SAT, RGM, RMA; Original draft: AK, SAT, RGM, and RMA; Review and editing: all authors; Funding acquisition: RGM and JLN; All authors have read concurred with the final version of the manuscript.

## Supplementary Figures

**Supplementary Figure S1:**
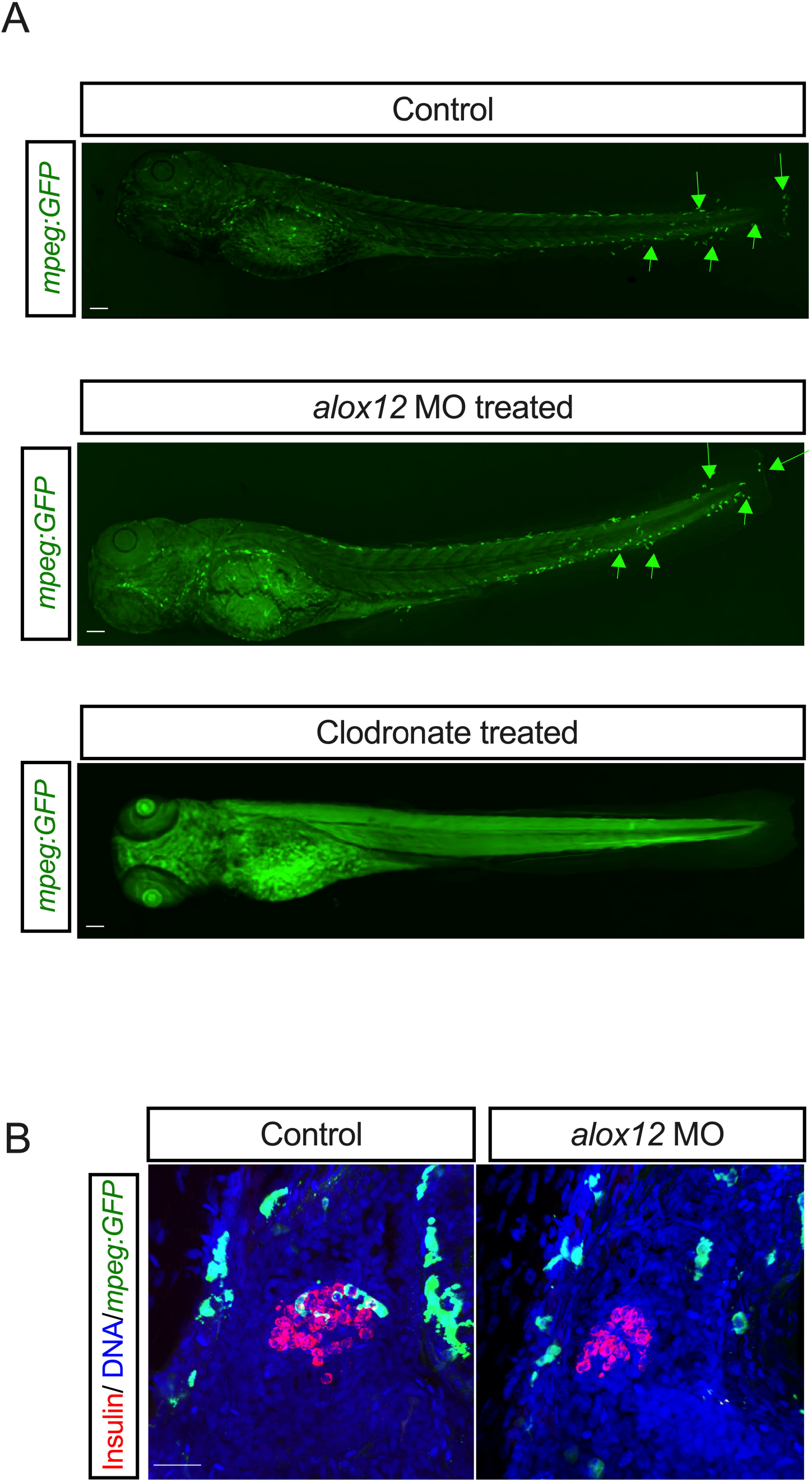
Effect of clodronate liposomes and *alox12* MO on whole-embryo macrophages. (***A***) Representative images of *Tg(mpeg:GFP)* zebrafish treated under control conditions, with *alox12* MO, or injected with clodronate liposomes and stained for GFP (macrophages, green). *Green* arrows indicate discrete macrophages; (B) Representative images of islets from *Tg(mpeg:GFP);Tg(ins:NTR)* zebrafish treated with MTZ, then stained for insulin (*β cells, red*), GFP (*macrophages, green*), and DAPI (*nuclei, blue*). Scale bars indicate 100 μm.

**Supplementary Figure S2:**
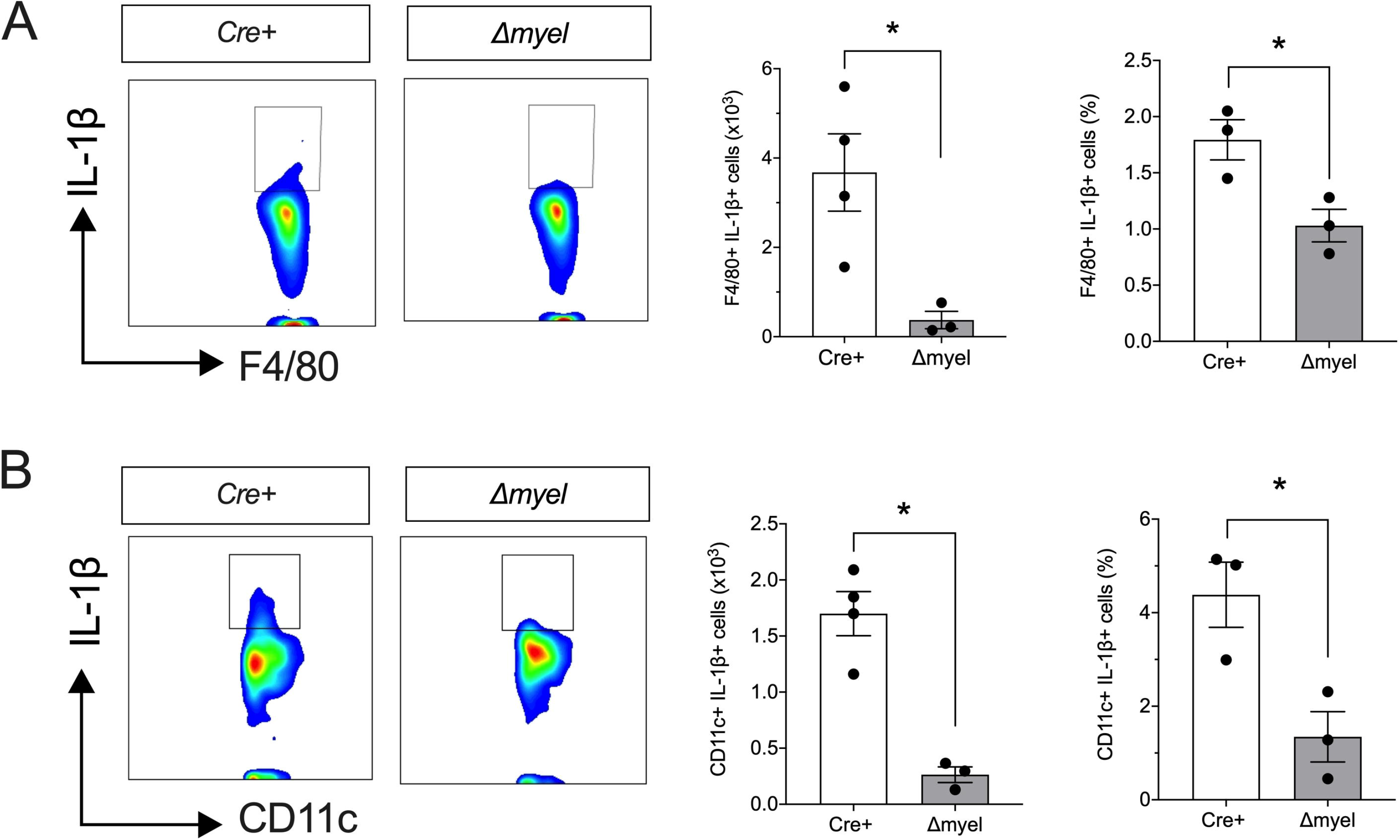
Reduced proinflammatory myeloid cell populations in pancreatic lymph nodes of *NOD:Alox15*^Δ*myel*^ mice. Pancreatic lymph nodes were isolated from *NOD:Lyz2-Cre* (*Cre*+) control mice and *NOD:Alox15*^Δ*myel*^ (Δ*myel*) mice at 8 weeks of age and subjected to flow cytometry analysis. (***A***) Representative contour plot showing gating of F4/80+ IL-1β+ cells (*left*), total number of F4/80+ IL-1β+ cells (*middle*), and F4/80+ IL-1β+ cells as a percentage of total cells (*right*); **(*B*)** Representative contour plot showing gating of CD11c+ IL-1β+ cells (*left*), total number of CD11c+ IL-1β+ cells (*middle*) and CD11c+ IL-1β+ cells as a percentage of total cells (*right*). All data are presented as mean ± SEM. \**P* <0.05 for the comparisons shown.

**Supplementary Figure S3:**
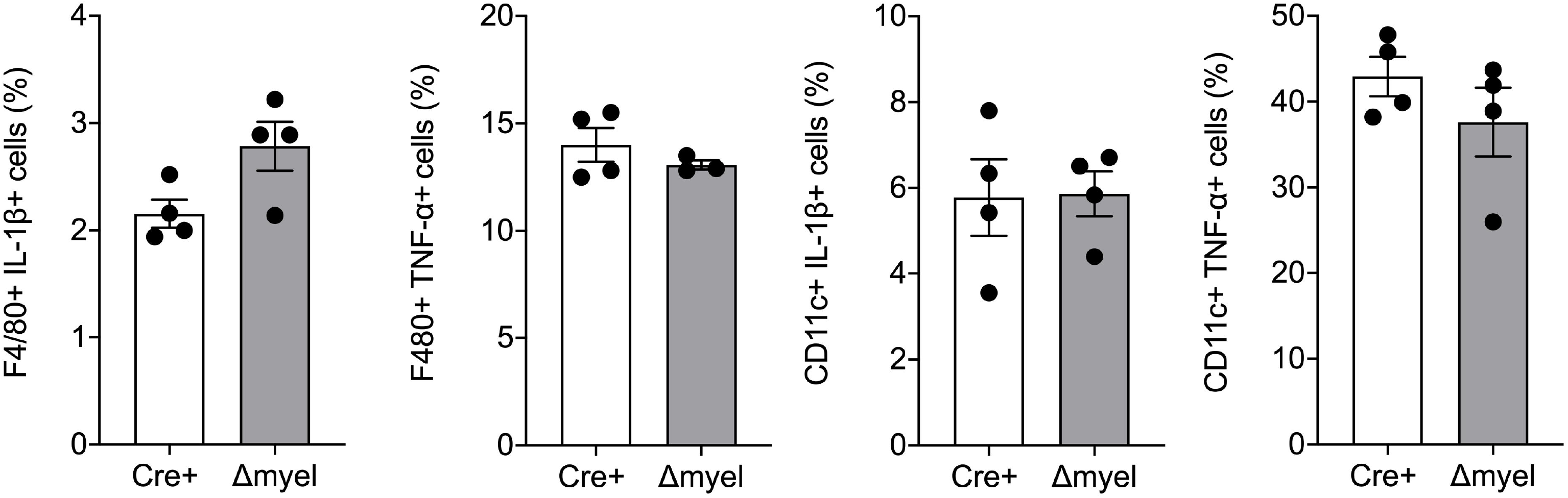
Unaltered myeloid cell populations in spleen of control *NOD-Lyz2-Cre* and *NOD:Alox15*^Δ*myel*^ mice. Spleens were isolated from control *NOD-Lyz2-Cre* (*Cre*+) and *NOD:Alox15*^Δ*myel*^ (Δ*myel*) mice at 8 weeks of age and subjected to flow cytometry analysis for proinflammatory myeloid cell populations. Shown are the F4/80+ IL-1β+ cells, F4/80+ TNFα+ cells, CD11c+ IL-1β+ cells, and CD11c+ TNFα+ cells as a percentage of total cells. Data are presented as mean□±□SEM.

**Supplementary Figure S4:**
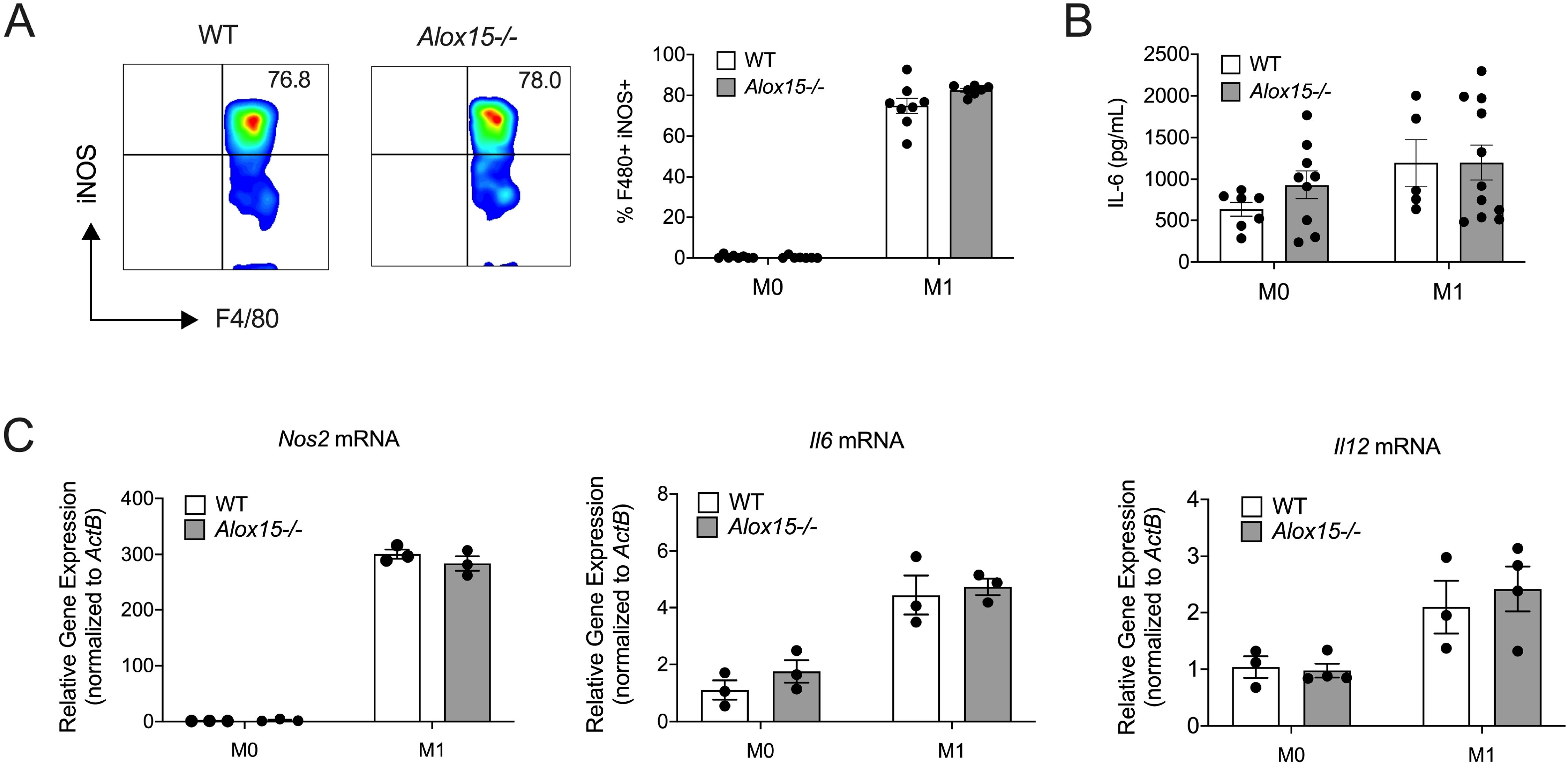
Effect of 12-LOX depletion on “M1” polarization of macrophages. Peritoneal cells were isolated from wildtype and *Alox15*-/- mice and unpolarized (M0) or polarized in vitro to a “M1-like” state upon incubation with lipopolysaccharide (LPS) and IFN-γ, then subjected to flow cytometry analysis and quantitative RT-PCR. (***A***) Representative contour plot showing gating of F4/80+ iNOS+ cells is shown on the *left* and quantitation of F4/80+ iNOS+ cells as a percentage of total cells is shown on the *right*; (***B***) IL6 levels in media of unpolarized (M0) and M1-polarized cells; (***C***) quantitative RT-PCR data for the indicated genes (normalized to *Actb*). All data are presented as mean□±□SEM. \**P* <0.05 for the comparisons shown.

**Supplementary Figure S5:**
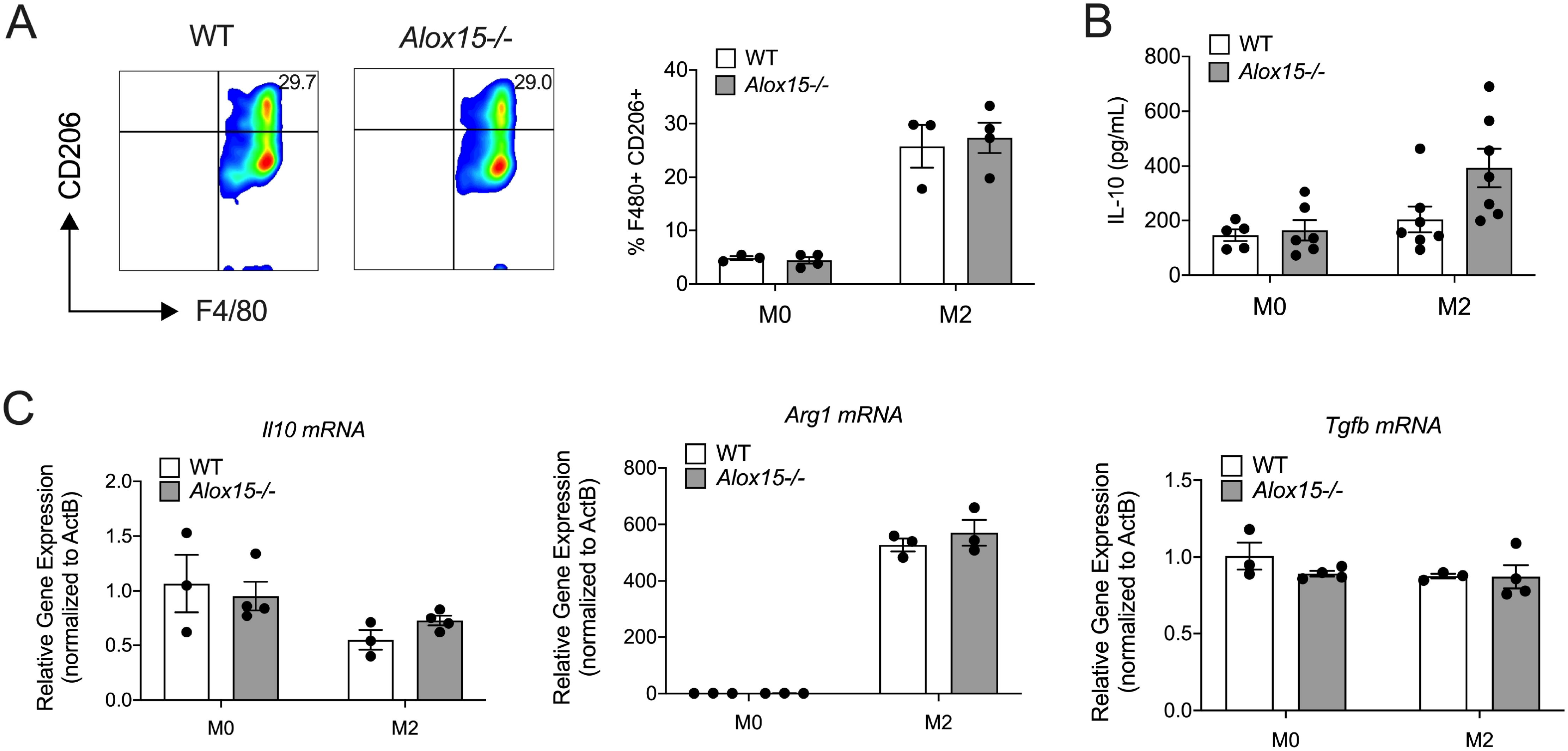
Effect of 12-LOX depletion on “M2” polarization of macrophages. Peritoneal cells were isolated from wildtype and *Alox15*-/- mice and unpolarized (M0) or polarized in vitro to a “M2-like” state upon incubation with IL-4, then subjected to flow cytometry analysis and quantitative RT-PCR. (***A***) Representative contour plot showing gating of F4/80+ CD206+ cells is shown on the *left* and quantitation of F4/80+ CD206+ cells as a percentage of total cells is shown on the *right*; (***B***) IL-10 levels in media of unpolarized (M0) and M2-polarized cells; (***C***) quantitative RT-PCR data for the indicated genes (normalized to *Actb*). All data are presented as mean□±□SEM. \**P* <0.05 for the comparisons shown.

